# Alpha satellite insertion close to an ancestral centromeric region

**DOI:** 10.1101/2021.03.10.434819

**Authors:** Giuliana Giannuzzi, Glennis A. Logsdon, Nicolas Chatron, Danny E. Miller, Julie Reversat, Katherine M. Munson, Kendra Hoekzema, Marie-Noëlle Bonnet-Dupeyron, Pierre-Antoine Rollat-Farnier, Carl A. Baker, Damien Sanlaville, Evan E. Eichler, Caroline Schluth-Bolard, Alexandre Reymond

## Abstract

Human centromeres are mainly composed of alpha satellite DNA hierarchically organized as higher-order repeats (HORs). Alpha satellite dynamics is shown by sequence homogenization in centromeric arrays and by its transfer to other centromeric locations, for example during the maturation of new centromeres. We identified during prenatal aneuploidy diagnosis by FISH a *de novo* insertion of alpha satellite DNA from the centromere of chromosome 18 (D18Z1) into cytoband 15q26. Although bound by CENP-B, this locus did not acquire centromeric functionality as demonstrated by lack of constriction and absence of CENP-A binding. The insertion was associated with a 2.8 kbp deletion and likely occurred in the paternal germline. The site was enriched in long terminal repeats (LTRs) and located ~10 Mbp from the location where a centromere was ancestrally seeded and became inactive in the common ancestor of humans and apes 20-25 million years ago. Long read mapping to the T2T-CHM13 human genome assembly revealed that the insertion derives from a specific region of chromosome 18 centromeric 12-mer HOR array in which the monomer size follows a regular pattern. The rearrangement did not directly disrupt any gene or predicted regulatory element and did not alter the methylation status of the surrounding region, consistent with the absence of phenotypic consequences in the carrier. This case demonstrates a likely rare but new class of structural variation that we name ‘alpha satellite insertion’. It also expands our knowledge on alphoid DNA dynamics and conveys the possibility that alphoid arrays can relocate near vestigial centromeric sites.

## Introduction

Alpha satellite is a class of highly repetitive DNA defined by a group of related, highly divergent AT-rich repeats or ‘monomers’, each approximately 171 bp in length. Alpha satellite, also named alphoid DNA, comprises up to 10% of the human genome and is mostly found tandemly repeated within constitutive heterochromatin at centromeres and pericentromeric regions. At centromeric regions, satellite monomers are hierarchically organized into larger repeating units, in which a defined number of monomers have been homogenized. These units, which are named ‘higher-order repeats’ (HORs), are tandemly arranged into chromosome-specific, megabase-sized satellite arrays with limited nucleotide differences between repeat copies ^1–6^.

The centromere is the chromosomal locus where sister chromatids attach and the kinetochore is assembled, which is essential for proper chromosome segregation during cell division. While alpha satellite DNA constitutes the sequence of all mature centromeres, it is not sufficient nor necessary for centromere identity. This is demonstrated by dicentric chromosomes that assemble the kinetochore at only one of two alpha-satellite regions ^7^ and analphoid chromosomes that possess fully functional centromeres ^8^. Centromere function appears to be epigenetically established and maintained by local enrichment of the CENP-A histone H3 variant within nucleosomes rather than presence of alphoid DNA ^9–12^. This function can be inactivated at an original site and moved to a new position along the chromosome ^13^. It is similarly turned off after a chromosomal fusion to ensure stability of the derived dicentric chromosome. These events determine the emergence of evolutionary new centromeres and the appearance of recognizable genomic regions where the centromere used to be positioned in the past ^14; 15^. Insights into the molecular steps of centromere repositioning from the birth of a new centromere to its maturity were uncovered by studying fly, primate, and equid chromosomes ^16; 17^. These analyses showed that new centromeres are first epigenetically specified and then mature by acquiring the satellite DNA array, in some cases going through intermediate configurations bearing DNA amplification ^18; 19^.

Besides the main pericentromeric and centromeric locations, smaller regions of alpha satellite DNA are located in the human genome >5 Mbp from the centromeres, with around 100 blocks annotated in the reference by the RepeatMasker program ^20; 21^. For example, three large blocks, respectively 11, 8, and 13 kbp long, are located within cytoband 2q21 with SVA (SINE/VNTR/Alu) and LINE elements intervening between them. These alphoid sequences are the relics of an ancestral centromere that became inactive ~5 Mya after the fusion of two ancestral chromosomes in the human lineage compared to big apes ^22–25^.

Here, we report an individual with a *de novo* insertion of an alpha satellite DNA array from the centromere of chromosome 18 into chromosome 15q26, the first observation of insertion of satellite DNA array outside centromeric and pericentromeric regions of the human genome that we are aware of. This case brings to light a probably rare and new class of structural variation and expands our knowledge on the spread and dynamics of alpha satellite.

## Results

### Prenatal, postnatal and family investigations

Amniocentesis was performed at 15 weeks’ gestation in a 35 years-old gravida 6 para 2 woman. She already had two healthy children, one miscarriage, and two pregnancies terminated due to fetal trisomy 21. Interphase FISH (Fluorescent *In Situ* Hybridization) on uncultured amniocytes with probes for main aneuploidies showed the presence of three signals for the alpha satellite DNA probe of chromosome 18 (D18Z1) in all cells (150/150) and three signals for chromosome 21 specific probes in 29 out of 121 cells (24%), suggesting a trisomy 18 and a mosaic trisomy 21. Karyotyping of cultured cells confirmed the presence of the mosaic trisomy 21 at 19% (12/62 cells) but showed the presence of two normal chromosomes 18. Metaphase FISH on cultured cells revealed the aberrant hybridization of the D18Z1 probe at chromosome 15q26 (**Figure 1A**). Chromosomal microarray did not show any imbalances, except the mosaic trisomy 21 (13%). FISH analysis of both parents showed that the alphoid DNA insertion was *de novo*. Pregnancy sonographic follow-up was normal. The proband, a healthy male baby, was born at term with normal birth parameters. Post-natal karyotype and FISH confirmed the mosaic trisomy 21 (6/33 cells; 18%) and the presence of the insertion of chromosome 18 alpha satellite on the long arm of a chromosome 15. At one year old, growth clinical examination (weight 10.6 kg, +1 standard deviation (SD); height 75 cm, +1 SD; occipito-frontal circumference 46.5 cm, +1 SD) and psychomotor development were normal, consistent with low level mosaic trisomy 21 and also suggesting that the alpha satellite insertion had no phenotypic consequences.

**Figure 1.**
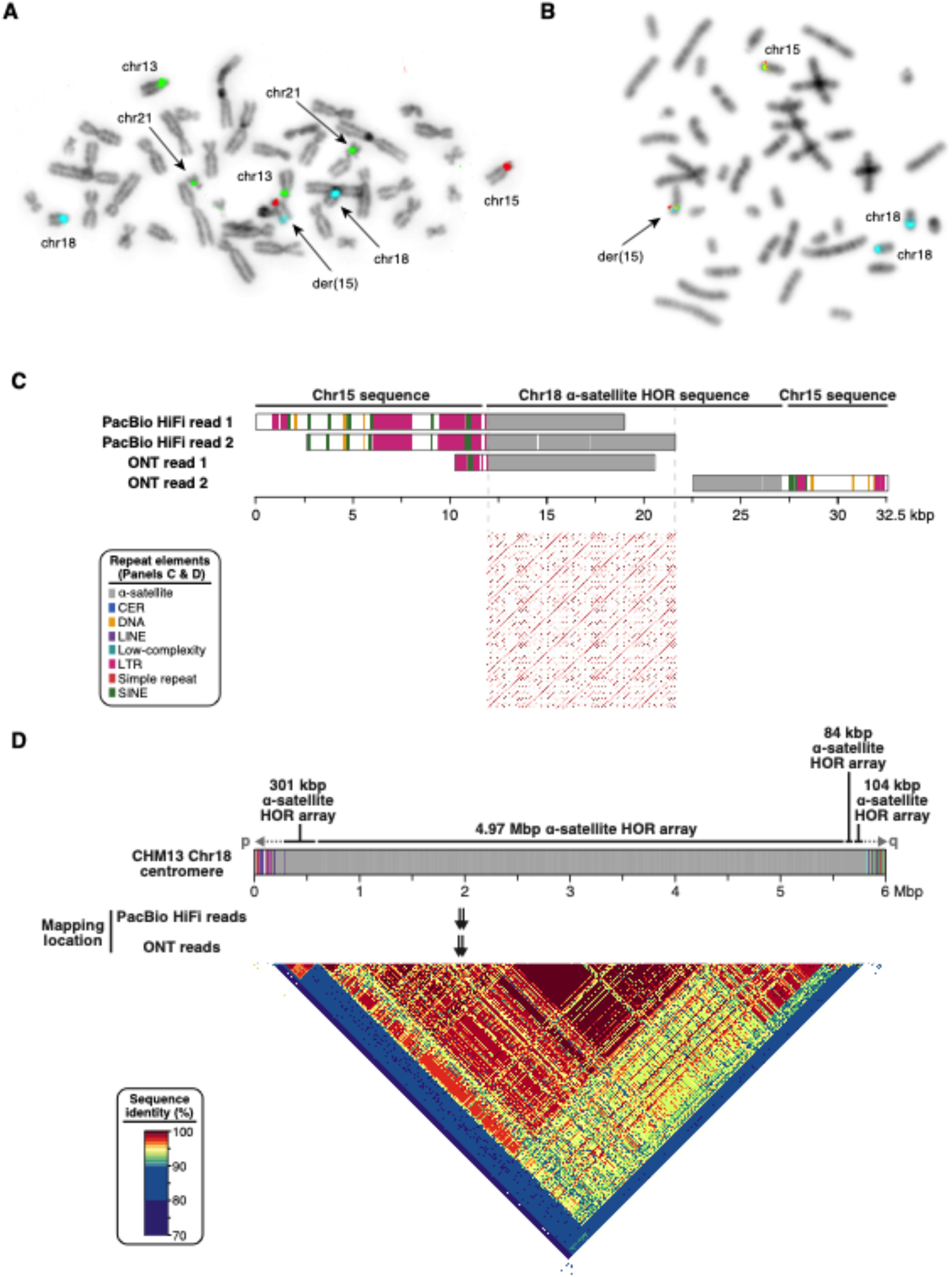
D18Z1 alpha satellite *de novo* insertion. **A)** FISH results of cultured amniocytes using alpha satellite DNA probes of chromosomes 15 (D15Z1, Texas-Red), 13/21 (D13/21Z1, green), and 18 (D18Z1, aqua) probes. **B)** FISH results of cultured amniocytes using the 15q25 BAC probes RP11-635O8 (red) and RP11-752G15 (green) flanking the ancestral centromere, and the D18Z1 (aqua) probe. **C)** Read length, repeat composition (color code in inset), and mapping location of the four selected HiFi and ONT reads (*top*). Dot plot (window size 20) of the 10 kb alpha satellite sequence from the centromere of chromosome 18 (*bottom*). **D**) Schematic representation of the CHM13-T2T chromosome 18 centromere with its repeat composition (*top*). A heatmap representation of sequence identity over the region is presented below. The mapping location of the PacBio HiFi and ONT reads is pinpointed by black arrows.

### Structural characterization of the rearrangement

To characterize the alphoid insertion at the sequence level, we performed WGS (whole genome sequencing) of the proband using the short-read Illumina platform. We first analyzed these data using a routine clinical analysis pipeline that did not identify any structural variant at chromosome 15q26. We then followed a customized approach, mapping reads to a library made up of the entire chromosome 15 and chromosome 18 centromeric alpha satellite DNA sequences. We isolated high-quality discordant paired reads mapped to both sequences, as well as chimeric reads anchored to chromosome 15 and containing alpha satellite DNA. These reads allowed us to define the positions of the proximal and distal breakpoints of the insertion at chr15:92,359,068 and chr15:92,361,920 (GRCh38), respectively. These coordinates, both subsequently validated by PCR, revealed the deletion of a 2,851 bp segment that was replaced by the insertion. We noted that the target site was ~10 Mbp distal from the position where an ancestral centromere was seeded and was shown to be active ~25 Mya in the common ancestor of Old World monkeys and apes, and was then inactivated sometime between 20 and 25 Mya in the common ancestor of the Hominoids (lesser apes, great apes, and humans) ^26^. This was further confirmed by the co-hybridization of the D18Z1 probe with two BAC probes flanking the ancestral centromere locus (RP11-752G15 and RP11-635O8) ^27^. This experiment showed, at metaphase resolution, that the satellite probe signal colocalized with both BAC probes on the derivative chromosome 15 (**Figure 1B**). The intensity and size of the FISH signal was similar to the ones of the BAC probes, suggesting that the inserted satellite DNA was ~50-300 kbp long.

We then sought to better characterize the rearrangement by generating long-read sequence information. We employed two technologies, ONT (Oxford Nanopore Technologies) with selective sampling via Read Until ^28^, targeting 50 kb of sequence on either side of the insertion, and PacBio HiFi sequencing. We sequenced the proband (~11.5x coverage at the targeted region), father (~20.1x), and mother (~19.8x) using readfish ^29^ on an ONT GridION, and the proband’s genome on one PacBio SMRT cell (~6.5x coverage). We confirmed the insertion breakpoints and the 2.8 kbp deletion but were unable to assemble a contiguous sequence spanning the entire insertion. To determine which parental chromosome the event occurred on, we phased the proband, father, and mother’s ONT reads and searched for diagnostic singlenucleotide variants that differed between the maternal and paternal haplotypes. The proband is hemizygous for two maternal variants mapping within the deleted region while the father is homozygous for the alternative allele. Conversely, the proband harbored one paternal variant on the haplotype with the insertion that is absent in his mother. This demonstrated that the rearrangement occurred on the paternal chromosome. Analysis of the junctions showed that, besides the aforementioned deletion, no further rearrangements, such as a target site duplication, occurred at the boundaries. At the proximal junction, a short sequence stretch of four nucleotides (CAAA) was identified that could not uniquely be assigned to the chromosome 15 or the satellite DNA. However, due to its small size, it is unlikely that this stretch of homologous sequence had a role in the rearrangement mechanism, particularly in the determination of the target site.

We analyzed the content of interspersed repeats in 5 kb segments upstream and downstream of the rearrangement breakpoints as well as in the deleted segment on chromosome 15 sequence. These segments were enriched for LTR (long terminal repeats derived from endogenous retroviruses) content when compared to the human genome average, as assessed by simulation for the entire 13 kb segment (4.34-fold, *P* = 0.035, **Table 1**).

**Table 1.** Content in interspersed repeat elements of the rearranged target site on chromosome 15. The “*E*” value is the enrichment coefficient that was calculated by dividing the observed value by the mean of 10,000 genome-wide permutations (human genome average).

|  | Sequence upstream of the insertion (5 kb) | Deletion (2851 bp) | Sequence downstream of the insertion (5 kb) | Entire region | Human genome average | *E*, *P* ± SE |
| --- | --- | --- | --- | --- | --- | --- |
| SINEs | 9% | 0% | 12% | 8% | 12% | 0.65, 0.57 ± 0.005 |
| LINEs | 0% | 0% | 0% | 0% | 19% | 0, 1 |
| LTR elements | 62% | 32% | 13% | 36% | 8% | 4.34, 0.035 ± 0.002 |
| DNA elements | 0% | 0% | 9% | 4% | 3% | 1.21, 0.3 ± 0.005 |

### Structural characterization of the alpha satellite DNA insertion

While we were unable to assemble the full sequence of the insertion, we investigated its structural properties by identifying reads with the longest content in alpha satellite DNA and unequivocally derived from this site, i.e. chimeric reads anchored to chromosome 15 sequence on either side of the insertion, spanning one breakpoint, and containing chromosome 18 centromeric alpha satellite sequences.

We selected two PacBio HiFi reads (PacBio HiFi read 1 and PacBio HiFi read 2) containing 7,199 and 9,821 bp of satellite DNA and having 99.95 and 99.48% estimated accuracy, respectively, both transitioning over the proximal junction; an ONT read with 8,618 bp of satellite DNA at the proximal junction (ONT read 1); an ONT read with 4,583 bp of satellite DNA at the distal junction (ONT read 2) (**Figure 1C**). Best alignments to the human genome reference (GRCh38) of alpha satellite segments from these four sequences showed identity with centromere reference models of chromosome 18 ^30; 31^. Alignments to the CHM13-T2T (Telomere-to-Telomere) genome ^5; 32^ resulted in unique locations for each read and pointed the origin of the insertion to a precise 10 kb region in the centromere of chromosome 18 (chr18:17500488-17510699) (**Figure 1D**). HiFi reads showed 99.6 and 99.1% identity with this region, respectively, with the remaining divergence not explained by sequencing errors (~0.4%) probably reflecting inter-individual differences in centromeric sequences. ONT reads showed lower identity values with the same region (94.5 and 94.4%, respectively), mainly due to errors in their sequence. As the estimated size of the inserted segment (order of hundreds kbp) is bigger than the size of the corresponding interval within chromosome 18 centromeric sequence, we hypothesize that this region is variable among humans and likely expanded in the proband or alternatively in his paternal lineage. Overall, these results confirmed that the insertion originated from chromosome 18 centromeric DNA and suggest that the CHM13 and our proband’s centromeres are structurally different in their sequence and size.

We next analyzed the repetitive structure of the satellite insertion using the corresponding T2T chromosome 18 centromeric sequence with ~99.99% accuracy. As chromosome 18 centromere is composed of two alpha satellite families, family I (D18Z1) and family II (D18Z2), both belonging to the suprachromosomal family 2 (SF2), whose arrays have a dimeric structure based on D1 and D2 monomers ^33^, we assessed the similarity with deposited sequences representing both families. Local pairwise alignments showed 98.8% and 81.7% identity, respectively with D18Z1 (M65181.1) and D18Z2 (M38466.1) sequences. Similar results, i.e. higher similarity with family I sequence, were obtained for both PacBio HiFi reads and the ONT read 2 transitioning over the distal breakpoint. These results indicate a closer relationship of the inserted satellite DNA to the D18Z1 family.

The T2T centromeric sequence showed a higher density of matches every ~2000 bp when assessed using the re-DOT-able tool (https://www.bioinformatics.babraham.ac.uk/projects/redotable/) (**Figure 1C**, *bottom panel*). To further assess this periodicity, we extracted 60 monomers, built a multiple sequence alignment, and visualized all pairwise identity percentages by creating two heatmaps. The first one shows monomers ordered according to their position in the array, while the second heatmap depicts monomers ordered according to the dendrogram determined by the hierarchical clustering of identity percentages (**Figure 2A**). In the dendrogram-based heatmap, the monomers cluster into two main clades formed by D1 and D2 monomers, as expected from the dimeric structure of the D18Z1 array. D1 and D2 monomers further group into 11 clades in agreement with their organization in a HOR unit of 12 monomers, with D1 monomers at positions 3 and 7 that are homogenized and form a single clade. D1 monomers at positions 9 and 11 and D2 monomers at positions 4 and 10 are also partly homogenized (**Figure 2A, right panel**). These results are consistent with a 12-mer HOR structure, matching the known organization of the D18Z1 satellite array ^2^. D1-D1/D2-D2 sequence identity ranges from 80.12 to 100% (median 85.96%); D1-D2 identity ranges from 64.67 to 77.19% (median 70.18%) (**Figure 2B**). While most monomers have a size of 171 bp, some D1 monomers are 166 or 167 bp long. The monomer size distribution is not random but follows a precise pattern in the HOR unit, showing another feature of the HOR hierarchical organization (**Figure 2C**) ^34^. Lastly, we grouped every 12 monomers into ~2 kb units and obtained four HOR repeats with 98.67-99.90% pairwise sequence identity (**Figure 2D**).

**Figure 2.**
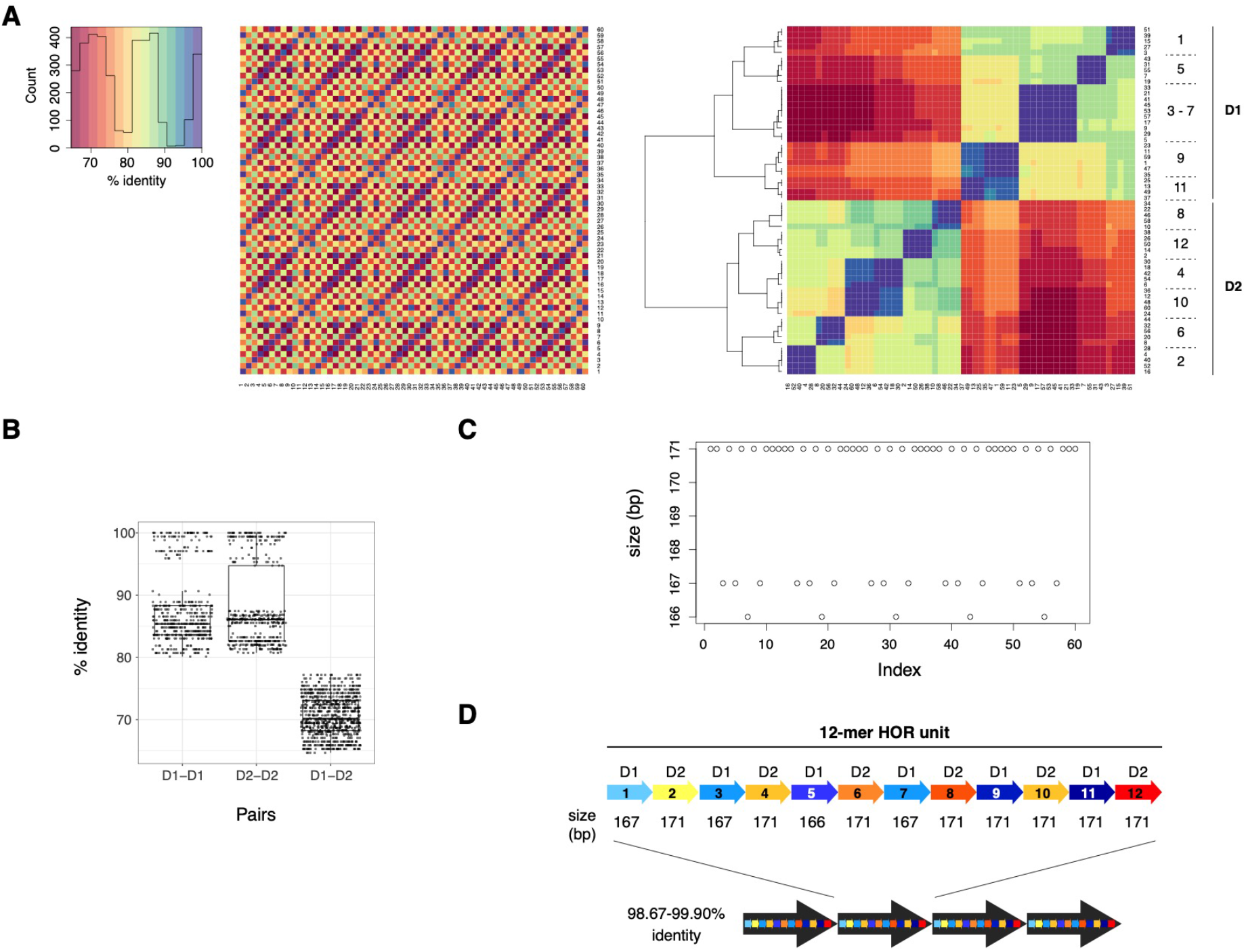
Organization of the alpha satellite array. **A)** Heatmaps of identity percentages between the 60 alpha satellite monomers derived from the T2T chr18:17500488-17510699 sequence, with monomers ordered either according to their position in the sequence (*left*) or as determined by clustering (*right*). In the latter, the position (1-12) in the 12-mer unit and monomer type (D1 or D2) are shown. **B)** Boxplots of identity percentages between D1-D1, D2-D2, and D1-D2 monomer pairs. **C)** Plot of monomer size with monomers ordered according to their position in the original sequence. **D)** Schematic of the 12-mer HOR units.

### Functional profiling of the rearranged site

To assess whether this structural change is likely to have functional impact, we examined gene annotation (GENCODE v32) at the insertion breakpoints as well as in the deleted region. We find that the rearrangement did not directly disrupt any gene, with the closest one *(ST8SIA2)* annotated 32 kb distally (**Figure 3A**). We then evaluated whether the rearrangement affected other functional elements, such as regulatory DNA. To this end, we leveraged publicly available data from the ENCODE consortium of chromatin activity measured by chromatin immunoprecipitation sequencing (ChIP-seq) for three histone modifications, i.e. methylated histone 3 at lysine 4 (H3K4me1), tri-methylated histone 3 at lysine 4 (H3K4me3), and acetylated histone 3 at lysine 27 (H3K27ac), on seven cell lines. These epigenetic marks are associated with poised enhancers (H3K4me1), promoters (H3K4me3), and active enhancers (H3K27ac). Neither the deleted segment nor the breakpoints overlapped any of these chromatin features, suggesting that the rearrangement did not disrupt a regulatory element (**Figure 3A**).

**Figure 3.**
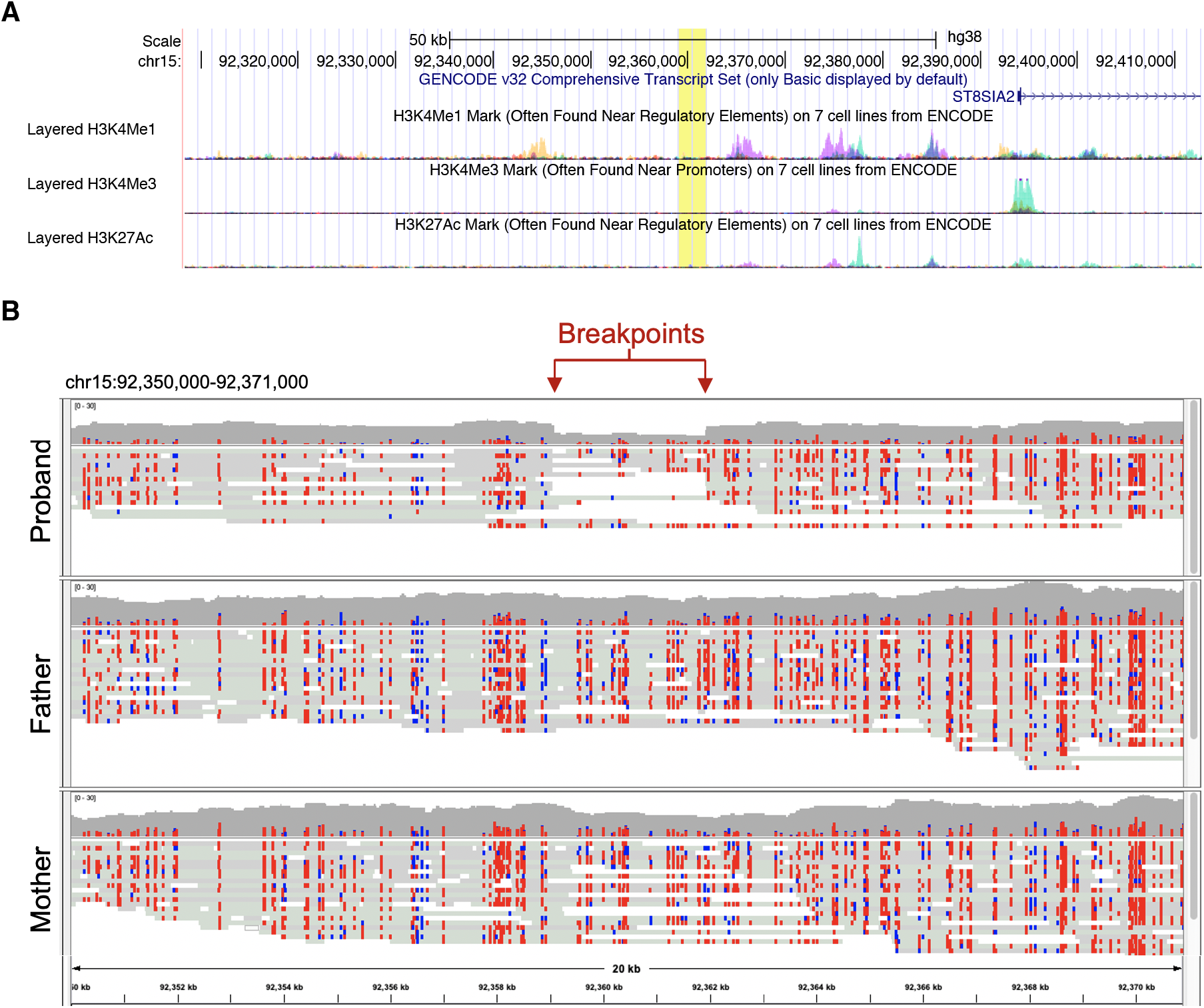
Functional profiling of the rearrangement site. **A) UCSC view of the 100 kbp region surrounding the rearrangement at 15q26.1**. The deleted region is highlighted in yellow, with deletion extremes corresponding to the satellite insertion positions. The GENCODE v32 and ENCODE regulations (H3K4me1, H3K4me3, and H3K27ac) tracks are shown (hg38). No gene and no enrichment of epigenetic marks found near regulatory elements are annotated in the deleted region. The closest gene, *ST8SIA2*, is mapped 32 kb distally. **B) Methylation pattern of the insertion site in the family trio**. Methylation data obtained from the ONT selective sequencing. Methylated (red) and unmethylated (blue) CpGs are shown. The methylation profiles are similar among the family trio.

Next, we assessed whether the insertion of centromeric satellite DNA, which comes from a heterochromatic locus, modifies the epigenetic status of the 15q26 target region. We leveraged CpG methylation data of the 20 kb genomic segment surrounding the insertion site using the ONT data of the proband and his parents. Cytosine methylation is an epigenetic modification often found in CpG dinucleotides that contributes to the formation of heterochromatic regions and leads to transcriptional modulation, in particular silencing. Comparison of the proband mutated allele with unrearranged ones, i.e. his maternal allele and the four alleles of his parents, revealed no major difference in the methylation patterns, indicating that the satellite insertion did not alter the methylation status of the surrounding region (**Figure 3B**). The absence of functional elements (gene or likely regulatory element) at the site and the maintenance of the methylation profile of the broader region suggest that the rearrangement itself has had no functional consequences. This is in line with the absence of clinical features in the proband that could not be explained by his trisomy 21 mosaicism.

### Immuno-FISH with anti CENP-A and CENP-B antibodies

Cytogenetic evaluation of the derived chromosome 15 revealed no chromosomal constriction at the position where the satellite DNA sequence was inserted, suggesting that this site did not acquire properties of a functional centromere. To further demonstrate this lack of epigenetically-defined centromeric function, we performed an immuno-FISH experiment with an antibody against the CENP-A protein. We observe no colocalization of the D18Z1 probe and CENP-A staining at the satellite insertion locus on the derivative chromosome 15 (**Figure 4**). We also assessed by immuno-FISH the binding of the CENP-B box by the CENP-B protein. In 20 out of 25 mitoses, we observe a faint pattern of staining of the CENP-B antibody corresponding to the satellite insertion, whereas in the remaining five we observed no signal (**Figure 4**). Such faint signals may derive either from the smaller size of the satellite insertion compared to a centromeric satellite array or to a weaker binding of the CENP-B protein. Nevertheless, these results suggest that CENP-B proteins recognize and bind the CENP-B box on the satellite monomers of the inserted sequence. Although CENP-B is not necessary and sufficient to confer centromeric function, it was shown that it creates epigenetic chromatin states permissive for CENP-A or heterochromatin assembly ^35^.

**Figure 4.**
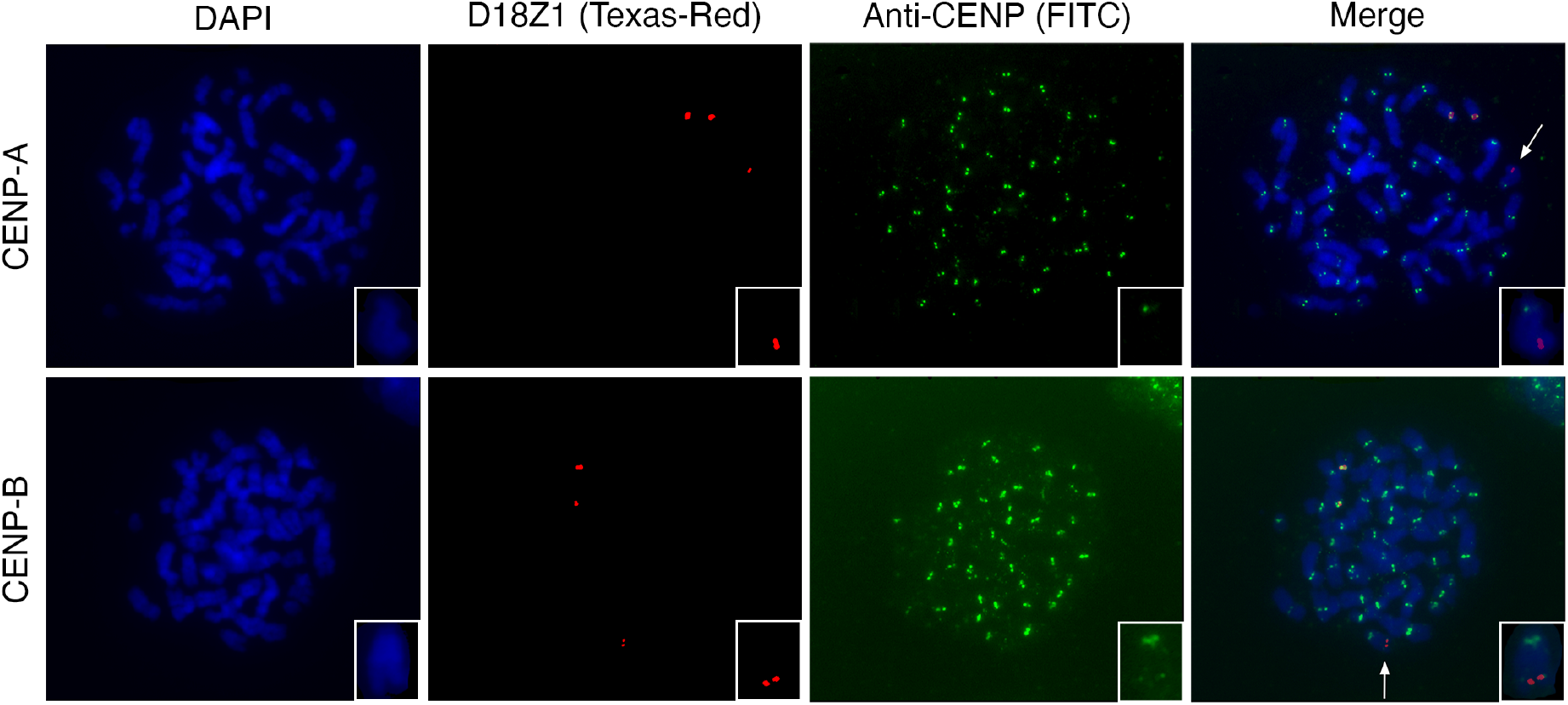
CENP-A and CENP-B immuno-FISH. Co-hybridization of the D18Z1 probe (red) with antibodies against CENP-A (*top*) and CENP-B (*bottom*) proteins (green) on chromosome metaphases from the proband. The arrows point at the derivative chromosome 15 that is also shown in larger magnification in the insets.

## Discussion

During routine prenatal testing for aneuploidy by FISH, we serendipitously identified an individual carrying a *de novo* insertion of alpha satellite DNA from the centromere of chromosome 18 into cytoband 15q26 (**Figure 5A**). Long-read sequencing and alignment to the CHM13-T2T genome showed that this segment originates from a precise location in the main chromosome 18 centromeric HOR array. Analysis of the repetitive structure showed novel features of chromosome 18 12-mer HOR, such as the presence of a regular pattern in monomer size, homogenization of D1 monomers at positions 3 and 7, partial homogenization of D1 monomers at positions 9 and 11, and partial homogenization of D2 monomers at positions 4 and 10.

**Figure 5.**
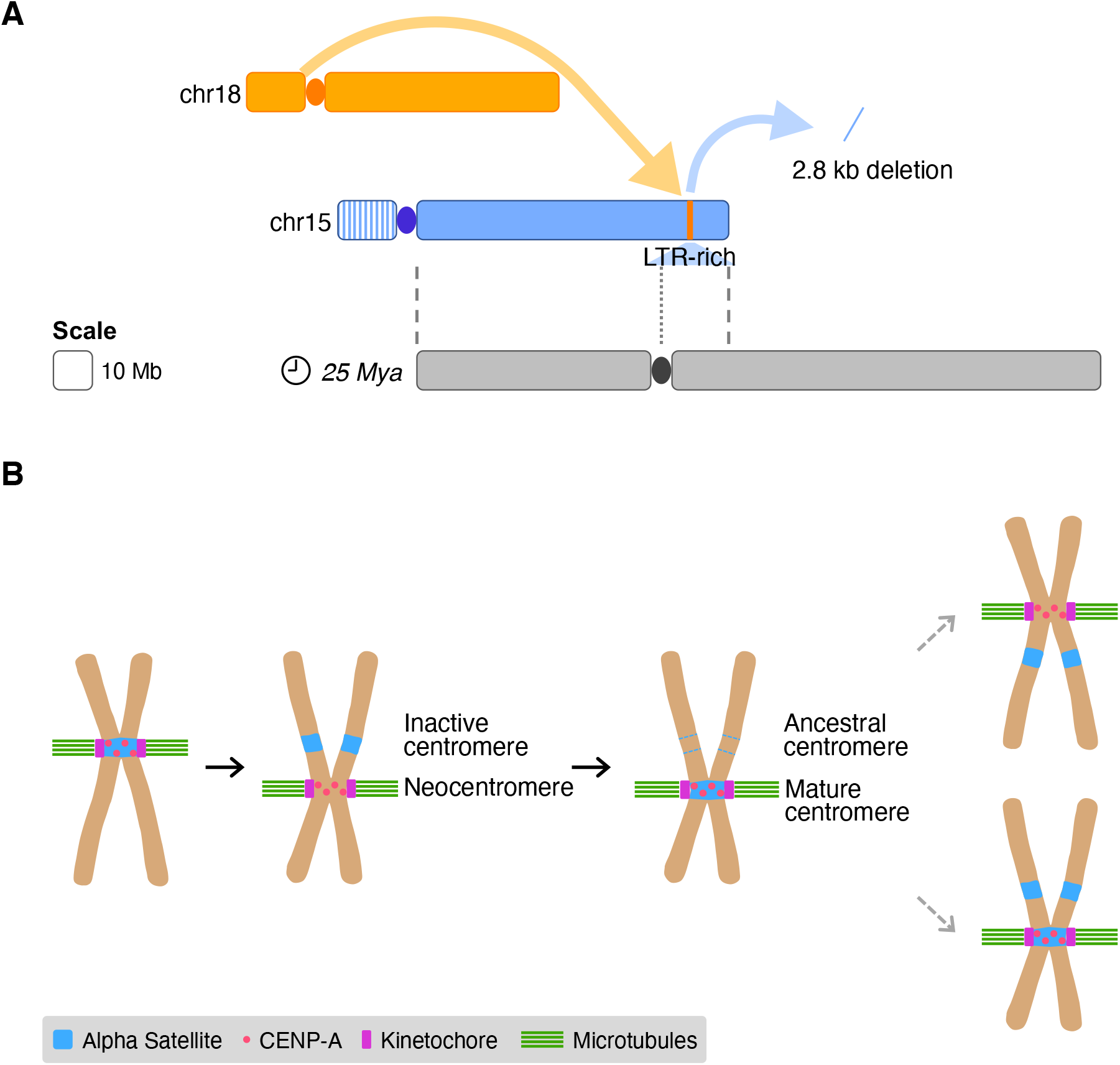
**A) Schematic overview of the rearrangement.** An alphoid array from the centromere of chromosome 18 inserted into an LTR-rich region of chromosome 15q26, ~10 Mbp distally from the site where an ancestral centromere was seeded ~25 Mya. This insertion was coupled with a 2.8 kbp deletion. Dashed lines pinpoint the boundaries of the synteny between chromosome 15 and the ancestral submetacentric chromosome; the dotted line indicates the position of the ancestral centromere. **B) Possible inferred model of alphoid DNA dynamics relative to centromere repositioning.** Following a centromere repositioning event, the new centromere is epigenetically specified by CENP-A binding and subsequently acquire alphoid DNA. It is possible that not only centromeric function but also the presence of alphoid DNA can be resurrected at ancestral centromeric sites.

Our report expands our knowledge on alpha satellite dynamics and proposes a new class of structural variation that we call ‘alpha satellite insertion’ (ASI). While our study describes an alphoid insertion into a non-centromeric/pericentromeric region, several prenatal FISH diagnostic reports describe the cross-hybridization of chromosome-specific centromeric alpha satellite probes to centromeric or pericentromeric regions of non-targeted chromosomes, i.e. the centromeres of chromosomes 19 and 22, the heterochromatin of chromosomes 1 and 9, and the pericentromeric region of chromosome 2 ^36–40^.

While the presence of some alphoid blocks outside centromeric and pericentromeric regions in the human genome reference ^20^ can be explained by the evolutionary history of the locus and past presence of a centromere, the existence of the others could result from fixed satellite insertion events. The maturation process of new centromeres that switch from satellite-free to satellite-rich regions (i.e. epigenetic specification followed by acquisition of the typical alpha satellite array), is telltale of the dynamism of alphoid DNA ^18; 19^. Novel localizations of alphoid DNA were reported in the white-cheeked gibbon, a lesser ape with an extensively rearranged karyotype when compared to the ancestral primate karyotype. In this species, alpha satellite DNA is found not only at centromeres but also at telomeres and interstitial positions corresponding to some evolutionary breakpoints ^41^.

Although the mechanisms governing alphoid insertion are not well understood, they are likely to involve a nonreciprocal transfer of DNA from a mature centromere via recombination, transposable elements, and/or rolling circle replication and reinsertion ^18; 42^. Our structural characterization of the rearrangement provides some insights on its mechanism of origin. The coordinated deletion suggests the involvement of double-strand breakage of DNA, as inferred for duplication events ^43^. It may be noteworthy that we also identified an enrichment of LTR elements in the long-range acceptor site. LTR retrotransposon activity is currently very limited or fully absent in humans ^44^ and therefore is unlikely to have directly driven the rearrangement. Such repeat-rich regions have been noted to be deleted as part of the duplication events associated with the new insertion of large (>100 kbp) blocks of segmental duplication ^43; 45^. Similarly, such coordinated deletions often (but not always) occur in gene-poor regions of the genome minimizing functional impacts of such massive new insertions and the fitness of the zygote/fetus.

The poor identification of alphoid DNA insertions outside centromeric regions until now could be linked either to the fact that they are extremely rare and/or because current sequencing-based methodologies and analytical approaches aimed at genotyping structural variants are opaque to these events due to their size and highly repetitive nature. Likewise, as only centromeric probes of chromosomes 18, X, and Y are routinely used to screen for aneuploidies prenatally, the ASI of other centromeric satellites and ASI smaller than the standard FISH resolution (~10 kb) could not be serendipitously found. While a designated analysis was performed to detect mobile element insertions (MEIs, including insertions of *Alu*, L1, and SVA) in 2,504 human genomes ^46^, the insertion of satellite DNA has not been specifically assessed in diverse human genomes. Similarly, our standard whole-genome sequencing diagnostic pipeline failed to identify the variant we describe.

Finally, the ASI location at 15q26 is noteworthy as a centromere resided at chromosome 15q25 in our past, ~10 Mbp away from the insertion site, and became inactive sometime between 20 and 25 Mya in the common ancestor of the ape lineage ^26; 27^. This feature raises the intriguing possibility that the alphoid array did not move to a random repeat-rich location in the genome, but instead revisited an evolutionary favored site mapping close to an ancestral centromere. Such a scenario together with the aforementioned prenatal reports proposes that alphoid DNA might preferentially move to other extant or past centromeric locations (**Figure 5B**). It also recalls previous studies that suggested that certain regions of the genome may have a “memory” and/or propensity to host centromeric function. As observed for the satellite insertion reported here, several analphoid clinical new centromeres seeded ~1–14 Mb from an ancestral centromeric site or a region that is orthologous to evolutionary new centromeres in other primate lineages ^26; 47–50^. This suggests that centromeric function and satellite array evolution may be restricted to region rather than precise chromosomal location.

The variant we identified hints that an alternative route to centromere formation might exist, where the region first acquires the satellite array and then the epigenetically-defined centromeric function emerges. Support for the latter comes from the observation that introduction of alpha satellite arrays in human cells can result in the formation of functional neocentromeres ^51; 52^.

Lastly, this case further highlights the risk of identifying false-positive aneuploidies of chromosomes 18, X, and Y when depending solely on centromeric satellite probes in rapid interphase FISH. Thus, it is critically important to follow up and confirm them by karyotyping.

## Material and Methods

### Short-read sequencing and data analysis

We extracted genomic DNA from cultured amniocytes of the proband using QIAamp DNA mini kit (Qiagen, Hilden, Germany). We performed 150 bp paired-end WGS using the short-read Illumina platform. We aligned the reads to the hg38 version of the human genome using BWA-MEM version 0.7.10 ^53^, run the BreakDancer version 1.4.5 ^54^ and ERDS version 1.1 ^55^, and visually inspected the 15q24-26 region using the IGV tool. As we identified no structural variant, we re-aligned the reads to a custom library made of chromosome 15 sequence (hg38) and a deposited sequence of alpha satellite family 1 of chromosome 18 (M65181.1) ^33^ using BWA version 0.7.17. To identify read pairs mapping at the insertion breakpoints, we selected discordant pairs with one end mapping on chromosome 15 and the other one on the satellite sequence and MAPQ>0. We removed soft and hard clipped reads and those mapping at the pericentromeric region of chromosome 15. We next identified chimeric reads spanning the breakpoints among the soft clipped reads using the Integrative Genomics Viewer (IGV) tool ^56^.

### Long-read sequencing and data analysis

We isolated PBMC (peripheral blood mononuclear cells) from the blood of the proband and both parents. We extracted DNA from approximately 1-2 million cells of actively growing culture by first pelleting the cells and resuspending them in 1.0 mL Cell Lysis Solution (Qiagen). The samples were incubated with RNase A solution at 37°C for 40 min. Protein Precipitation Solution (Qiagen) was added at 0.33x and mixed well. After a 10 min incubation on ice, the precipitate was pelleted (3 min, 15000 rpm, 4°C). The supernatant was transferred to new tubes, and DNA was precipitated with an equal volume of isopropanol. The DNA was pelleted (2 min, 15000 rpm, 4°C) and the pellet was washed three times with 70% EtOH. The clean DNA was rehydrated with DNA Hydration Solution (Qiagen) and left for two days to resuspend.

We generated a PacBio HiFi library from the proband’s genomic DNA using g-TUBE shearing (Covaris) and the Express Library Prep Kit v2 (PacBio), size selecting on the SageELF platform (Sage Science) to give a tight fraction of around 23 kbp by FEMTO Pulse analysis (Agilent). The library was sequenced on one SMRT Cell 8M using v2 chemistry, and we obtained 20.5 Gbp of HiFi reads with mean length of 20.9 kbp and median quality of Q27. We assembled the data with HiCanu ^57^ and Hifiasm ^58^ and aligned reads to the GRCh38 (hg38) reference genome using pbmm2 (https://github.com/PacificBiosciences/pbmm2).

Adaptive sampling was performed on an ONT GridION (one flow cell per sample) using readfish ^29^. For each sample 1.5 ug of DNA was used to prepare a LSK-109 library according to the manufactures protocol. DNA was sheared in a Covaris g-TUBE at 6 k rpm for 2 min. The region targeted was chr15:92,309,068-92,411,920 (hg38 coordinates). ONT FAST5 files were base-called using guppy 4.0.11 using the high-accuracy model. FASTQ files were pooled and aligned to hg38.no_alt.fa using both minimap2 ^59^ and ngmlr ^60^. We identified reads spanning the breakpoints (located at chr15:92,359,068 and chr15:92,361,920) by manual inspection of the 15q26 read alignments in IGV v2.4.16 ^56^. We called and phased variants using Longshot ^61^ and called CpG methylation using Nanopolish ^62^. Selected PacBio and ONT reads were aligned to the CHM13-T2T genome using pbmm2 and minimap2, respectively.

### Analysis of repeat element content

We assessed the content in repeat elements in the deleted segment and in the 5 kb segments upstream and downstream the insertion breakpoints by using the annotation of the GRCh38 RepeatMasker track ^63^. The null distributions were generated by performing 10,000 permutations of the entire 12,851 bp segment, excluding gaps and centromeres, by using BEDTools version v2.30.0 ^64^. R v4.0.3 ^65^ was used to compute empirical *P* values. Standard error (SE) was estimated using the formula SE=sqrt(*P*\*(1-*P*)/10,000).

### Satellite monomer and HOR analysis

Dot plots were created using the re-DOT-able tool (https://www.bioinformatics.babraham.ac.uk/projects/redotable/). We extracted satellite monomers by blast alignment ^66^ with D1 monomer sequence (AJ130751.1). We performed multiple sequence alignments of monomers using Muscle ^67^ with default options. We created heatmaps and plots using the gplots v3.1.0 (https://CRAN.R-project.org/package=gplots) and ggplot2 v.2.2.1 ^68^ packages in the R software environment ^65^.

### FISH and Immuno-FISH

FISH on uncultured amniocytes was performed with the Aquarius FAST FISH Prenatal kit (Cytocell, Cambridge, UK) (DXZ1, DYZ3, D18Z1, *RB1, DYRK1A* probes) according to manufacturer’s instructions. Metaphase spreads were prepared from amniotic fluid cells and lymphocytes according to standard procedures. FISH was further performed using BAC probes localized in 15q25.2, RP11-752G15 (FITC) (chr15:82,627,211-82,802,988, hg38) and RP11-635O8 (chr15:82,023,617-82,178,139) (TRITC) (RainbowFish, Empire Genomics, Buffalo, New York, USA) and alpha-satellites probes for chromosomes 15 (D15Z1, Texas-Red), 18 (D18Z1, Aqua) and 13/21 (D13/21Z1, Green) (Cytocell).

Immuno-FISH was performed on lymphoblastoid cells from the patient. Metaphase cells spreads were prepared according to a protocol adapted from Jeppesen ^69^. Briefly, lymphoblastoid cells were harvested after 44 hour culture, incubated at 37°C with colchicine (0.2μg/mL final concentration) during 2 hours, then in a 75 mM KCl hypotonic solution during 25 min. After centrifugation, cell pellet was resuspended in 75mM KCl/0.1% Tween20 and then cytocentrifuged 5 min at 1000 rpm. The slides were transferred to a Coplin jar containing KCMc solution (120 mM KCl, 20 mM NaCl, 10 mM Tris-HCl, pH 7.5, 0.5 mM EDTA, 0.1% (v/v) Triton X-100) and incubated 15 min at room temperature. Then immuno-FISH was performed with a protocol derived from Solovei et al. ^70^ using as primary antibodies mouse anti-CENP-A (Abcam, Ab13939) (1/200) and mouse anti-CENP-B (5E6C1 clone, generous gift from Hiroshi Masumoto, Japan) (1/200); AlexaFluor conjugated goat anti-mouse as secondary antibody (1/1000) and D18Z1 probe (Texas-Red) (Cytocell). Images were performed with a Zeiss AxioImager Z2 fluorescence microscope equipped with a CoolCube Camera.

## Acknowledgements

GG is recipient of a Pro-Women Scholarship from the Faculty of Biology and Medicine, University of Lausanne. This work was supported by the Swiss National Science Foundation grant 31003A_182632 and the Jérôme Lejeune Foundation to AR, by the National Institutes of Health grant HG010169 to EEE, and a grant from the Brotman Baty Institute for Precision Medicine to DEM and EEE. We wish to thank Emilie Chopin (Cell Biotechnology Center, Hospices Civils de Lyon) for providing lymphoblastoid cell lines, as well as Patrick Lomonte for his kind gift of antibodies and his advice.

